# Chronic stress remodels gut bacterial ecology and polysaccharide metabolism with neuroendocrine consequences despite unchanged behavior

**DOI:** 10.64898/2026.10.09.757984

**Authors:** Amelia C. Slusser, Tulasi Rao Relangi, LaRae A. Hudson, Magdalene M. Hernandez, Danny T. Le, Tyler J. Earp, Aaron T. Wright

**Affiliations:** Department of Biology, Baylor University, Waco TX 76708, United States

## Abstract

The gut-microbiome-brain axis (GMBA) links the gut and brain through metabolic, immunological, and neuroendocrine pathways, and can become dysregulated during psychological stress (PS). While previous studies primarily infer microbial function and use physical stressors that incompletely model human PS, here, functional multi-omics was applied to a chronic social-isolation mouse model. Although isolation caused no measurable behavioral changes, brain proteomics revealed stress-associated molecular responses. Taxonomic and short-chain fatty acid analyses suggested limited microbiome changes; however, metaproteomics uncovered sex-specific restructuring of bacterial ecology and carbohydrate-active enzymes. In females, isolation increased *Petralouisia muris* proteins involved in bacterial microcompartment metabolism and β-glucuronidases across *Lachnospiraceae* taxa. Enriched β-glucuronidase abundance coincided with reduced circulating glucuronides and altered brain estrogen-receptor activity, suggesting a microbial deconjugation-host estrogen-signaling interaction. These findings demonstrate that functional GMBA alterations may precede or occur independently of overt behavioral phenotypes and highlight the value of direct functional profiling in PS.

## Introduction

Psychological stress (PS) is pervasive in modern life, affecting individuals across socioeconomic classes and cultural contexts. Young adults (18-34) in the United States report an average stress level of 6/10 (where 1 is “little to no stress” and 10 is a “great deal of stress”), and these levels have risen steadily over the past five years^1^. Although PS is adaptive and can even be beneficial over brief periods^2^, chronic stress increases the risk for developing cardiovascular diseases^3^, metabolic syndromes^4^, psychiatric disorders^5,6^, and even some neurodegenerative diseases^7,8^. Beyond the brain and the endocrine system, chronic PS also disrupts peripheral systems like the gastrointestinal tract and can predispose individuals to diseases such as irritable bowel syndrome and inflammatory bowel disease^9,10^.

Communication between the gut and the brain occurs through the gut-microbiome-brain axis (GMBA), a complex, bidirectional network links the brain to the gut through a variety of overlapping metabolic, immunological, and neuroendocrinal pathways^11–13^. Signals from the host have been shown to shape microbial ecology, gut motility, and mucosal immunity while microbial activities are known to regulate neural plasticity, cognition, stress reactivity, and blood-brain barrier permeability^11,12,14^. Thus, the gut microbiome has been recognized not as a passive target, but as an active participant in shaping host responses to PS.

The GMBA is a delicate interplay that is easily dysregulated by stress, resulting in profound effects on the gut. Growing evidence indicates that chronic PS instills inflammation in the intestine, increases gut permeability and disrupts homeostasis^15^. For example, recent work demonstrated that stress-induced enrichment of *Lactobacillus murinus* increased metabolism of tryptophan into indole-3-acetate, triggering epithelial injury through intestinal inflammation^16^. Other research links PS to shifts in microbial community composition^17,18^. Although consensus on key species remains limited, studies suggest that PS creates an intestinal environment favoring pro-inflammatory species such as mucin degraders and enteric pathogens while depleting beneficial taxa like *Lactobacillus, Faecalibacterium,* and *Bifidobacterium*^18–20^.

Despite extensive research, key gaps remain in GMBA studies. Much of the field relies on correlations between host behavior, stress exposure, and microbial taxonomy. While valuable for identifying broad community shifts, these approaches assume that changes in composition reflect changes in function. However, microbial taxa exhibit substantial functional redundancy^21^, and closely related species may differ in expressed metabolic activity depending on environmental context^22,23^. Advances in next-generation sequencing and metabolomics have shifted the emphasis toward predicted functional changes^18^. Yet, metagenomic predictions reflect only genetic potential and may not capture true metabolic activity. Metabolomics can measure actual metabolic output, but without genomic or proteomic context, such data are challenging to integrate and interpret. Furthermore, metabolomic reference databases remain poorly annotated, and many detected features remain unidentifiable, limiting discovery of novel GMBA mechanisms^24^.

Another challenge in the GMBA field is the translational relevance of rodent stress models. Many experimental models rely on genetic manipulation, environmental or physical stressors, or chemical injections to induce a behavioral response^25^. While useful for generating robust phenotypes, these approaches do not always reflect the nature of stress experienced by humans^26^. In humans, chronic stress and associated disorders generally arise from psychological or social stress rather than physical or environmental stressors^27,28^. Therefore, a social isolation (SI) model represents a biologically relevant stressor that may better bridge rodent models and the human experience. Despite translational relevance, the impact of SI stress on the functional activity of the gut microbiome and associated host phenotypes remains poorly understood.

In this study, a multi-omic approach integrating taxonomic, metaproteomic, host proteomic, and metabolomic measurements was used to functionally investigate the effects of SI stress on the GMBA. A murine SI paradigm was designed to model a biologically relevant form of chronic PS. Although the model did not produce robust behavioral differences, host proteomics revealed molecular changes consistent with stress-associated phenotypes. To our knowledge, this is the first study to functionally characterize the GMBA in a murine model of chronic stress without measurable behavioral changes. Taxonomy-based analysis suggested that SI stress had minimal effects on gut microbial community composition; however, metaproteomics revealed functional changes in microbial stress response pathways, altered bacterial community ecology favoring propanediol-metabolizing *Petralouisia muris*, and community-wide restructuring of carbohydrate metabolism. In females, changes in microbial function were accompanied by changes in circulating glucuronide candidates and brain proteomic profiles consistent with dysregulated estrogen signaling. Together, these findings provide a window into how SI-induced stress can produce functionally meaningful changes to the GMBA without robust behavioral or taxonomic alterations. Broadly, this work highlights the importance of moving beyond inference-based analyses toward direct, functional measurements to better understand the GMBA in translational models of chronic PS.

## Results

### Social isolation resulted in a stress-response phenotype at the molecular level

Adult male (n=6) and female (n=6) C57BL/6J mice were subjected to SI for 6 weeks to evaluate the effects of chronic PS on the metabolic profile of gut microbiota and associated brain functioning (Fig. 1A). To reduce visual interaction among isolated animals, isolation cages were wrapped in masking tape (Fig. 1B). Over the 6-week treatment period, SI mice did not exhibit significant changes in body weight or behavioral testing (FST) compared to group-housed (GH) controls (ns; S1; Fig. S1A-C). This suggests the model used in this study did not result in significant behavioral changes due to SI-induced stress.

**Figure 1.**
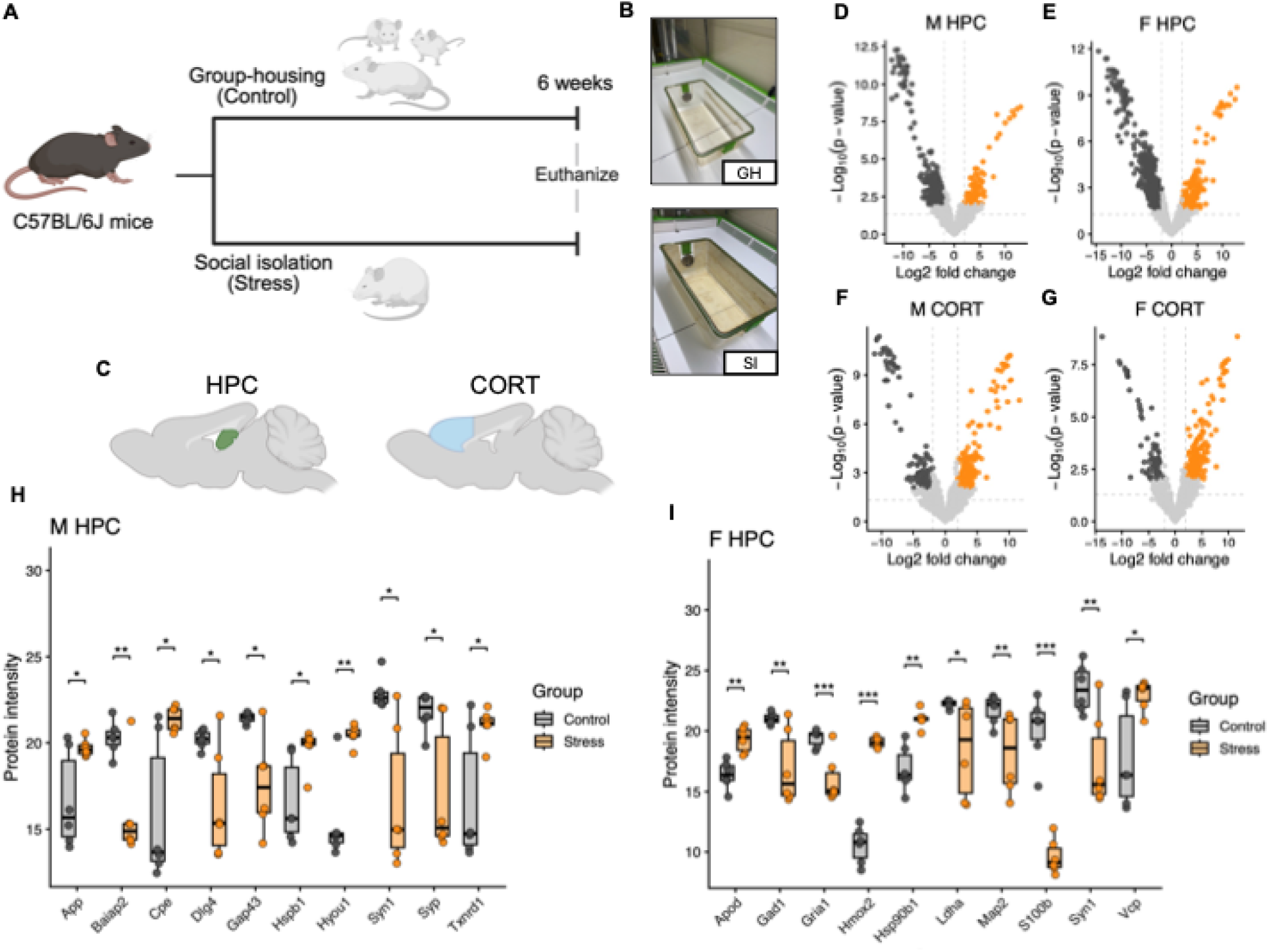
Social isolation resulted in a molecular stress-response phenotype, but no measurable behavioral changes. **A** Six-week isolation stress model. A total of 24 mice were included in this study: Male GH n=6, male SI n=6, female GH n=6, female SI n=6. **B** GH and SI cage design. **C** Brain regions sampled for protein profiling to determine molecular effects of stress. **D-G** Volcano plots demonstrating differential expression of proteins in the male (M) and female (F) hippocampus (HPC) and cortex (CORT). **H & I** Expression of HPC proteins dysregulated after the 6-week isolation treatment compared between control and stress in males and females. Statistical significance was calculated using a weighted linear model and empirical-Bayes moderated t-test method through the limma R package (version 3.66.0). Raw *p*-values were adjusted using Benjamini-Hochberg (BH) false discovery rate (FDR) adjustment (*q*-value). Significance was defined as *q*<0.05 and an absolute Log2(fold change) ≥ 2: \**q*<0.05, \*\**q*<0.01, \*\*\**q*<0.001. Whiskers extend to the most extreme values within 1.5X the interquartile range.

Given the absence of significant behavioral differences in this cohort, global proteomic analyses were performed on the hippocampus and cortex, two brain regions that are highly sensitive to PS, using DDA LC-MS/MS (Fig. S1D-S; Fig. 1C). Across hippocampal samples, 5,253 proteins were detected in males, and 5,024 proteins were detected in females. In the cortex, 6,045 proteins were detected in males, and 6,145 proteins were detected in females (S2 & S3).

Filtering and statistical analysis revealed widespread stress-associated proteomic changes in the hippocampus (Fig. 1D & E). In males, 221 proteins that were significantly decreased and 83 significantly increased in the SI mice compared to the GH controls (*q*<0.05, *abs(log_2_(fc))*>2; S2; Fig. 1D). In females 442 proteins significantly decreased and 122 significantly increased following SI (*q*<0.05, *abs(log_2_(fc))*>2; S2; Fig. 1E). Among significantly altered hippocampal proteins, 36 were detected only in male GH mice, 9 were detected only in male SI mice, 64 were detected only in female GH mice, and 17 were detected only female SI mice (S2).

Stress-associated proteomic differences were also observed in the cortex following filtering and statistical analysis (S3; Fig. 1F & G). In males, 103 proteins were significantly decreased, and 192 proteins were significantly increased in SI mice, while females displayed 68 significantly decreased and 167 significantly increased cortical proteins following the SI stress model (*q*<0.05, *abs(log_2_(fc))*>2; S3; Fig 1F & G). Among significantly altered cortical proteins, 31 were detected only in male GH mice, 24 were detected only in male SI mice, 16 were detected only in female GH mice, and 18 were detected only in female SI mice (S3). Taken together, these data suggest that the SI stress model produced drastic region- and sex-specific shifts in the protein profile of the brain.

Several of the significantly altered hippocampal proteins were associated with stress response and proteostasis (*Hspb1, Hyou1, Hsp90b1, Txnrd1, Vcp, Hmox2, Apod*) synaptic structure and scaffolding (*Dlg4, Baiap2, Map2, Gap43*), signal transduction and neurotransmission (*Syn1, Syp, Gad1, Gria1*), neural protection and glial signaling (*S100b, Apod, App*), and metabolic or peptide-processing pathways (*Ldha, Cpe*; Fig. 1H & I). As these pathways are canonical indicators of a stress response^29–32^, and the abundance and activity of the specific proteins are known to change due to stress and neurological disorders^33–49^, the model used here was sufficient to induce detectable stress-associated molecular remodeling in the brain. Additionally, epithelial immune signaling and tight junction proteins in the colon were significantly disrupted in both male and female SI mice (*p*<0.05; S4; Fig. S1T & U), indicating that stress-associated molecular changes extended beyond the brain to the intestinal environment.

### Conventional methods to investigate the gut brain axis suggest minimal changes to the gut microbiome

To evaluate whether SI stress altered the gut community composition, 16S rRNA gene sequencing was performed on intestinal contents (pooled cecum and colon; S5; Fig S2A & B; Fig. 2A-C). Alpha diversity did not differ significantly between SI and GH mice in either males or females (ns; S5; Fig. S2C & D). Beta diversity was unchanged in female control mice compared to isolated mice (*p*=0.1126; S5; Fig. 2E), but male SI mice exhibited a significant shift in community structure compared to the male GH controls (*p*=0.0051; S5; Fig. 2D). Differential abundance analysis with MaAsLin2^50^ identified no significantly altered microbial species in females (ns; S5), but uncovered a significant decrease in *Roseburia zhanii* and *Roseburia faecis* in SI male mice (*q*<0.05; S5; Fig. S5E & F). These results indicate that the SI stress model produced limited detectable changes in community composition using 16S rRNA gene-based profiling.

**Figure 2.**
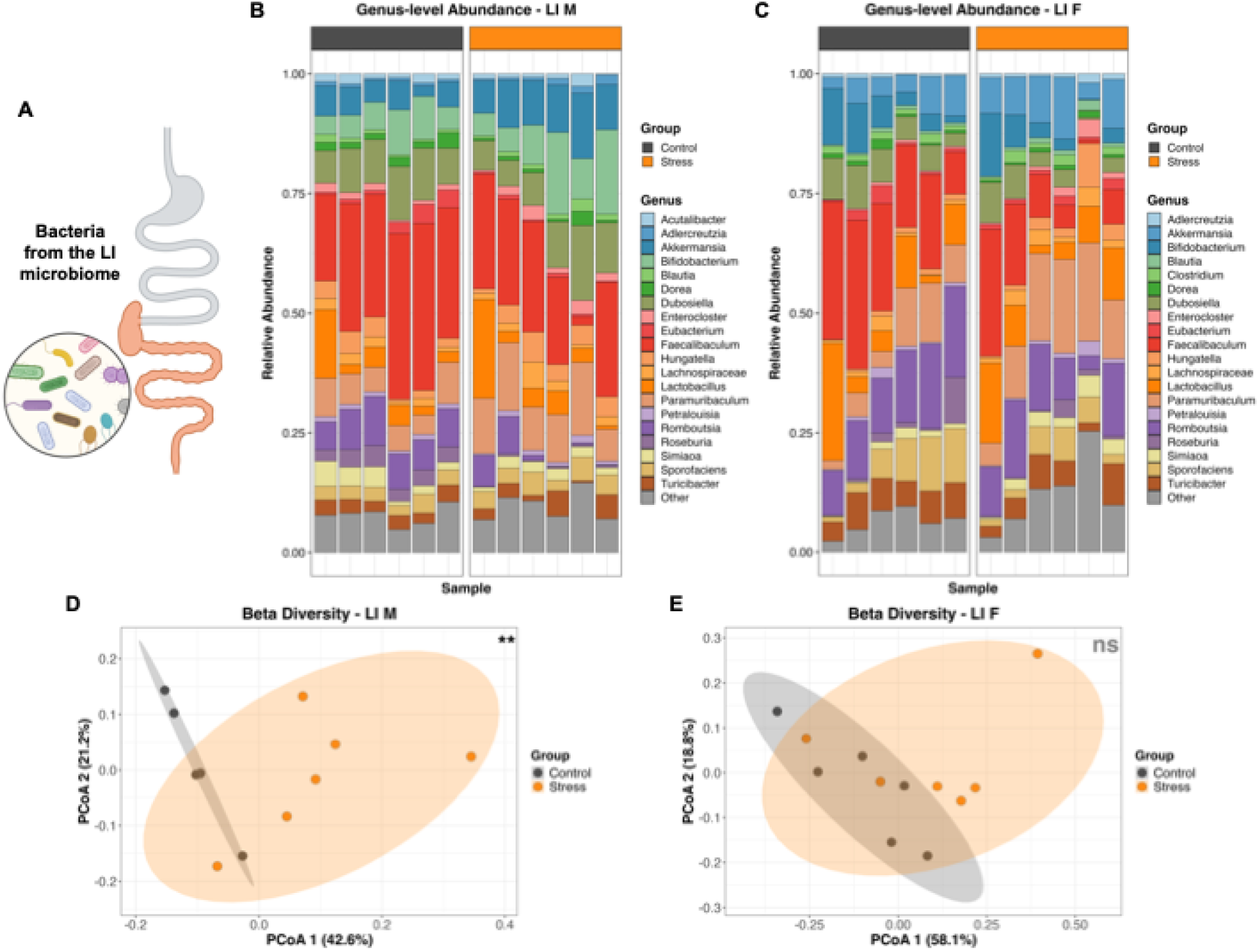
Conventional methods to investigate the gut-brain-axis suggest minimal changes to the gut microbiome. **A** Bacterial samples were collected from the cecum and colon and pooled for analyses (n=6, per group). **B & C** Stacked bar graphs showing relative abundance of bacterial genera composition in the large intestine of control and stressed mice, male and female, by sample replicate. **D & E** Beta diversity (Bray-Curtis Dissimilarity) PCoA plots of male and female large intestine microbiota. Statistical differences were calculated using permutational multivariate analysis of variance (PERMANOVA; adonis2, 9,999 permutations). Significant effect size was determined as *p*<0.5: not significant (ns), \**p*<0.05, \*\**p*<0.01.

Targeted metabolomics analysis did not reveal any significant differences in acetate, propionate, or butyrate between GH and SI mice (ns; S6; Fig. S2G & H). Taken together, conventional compositional and targeted SCFA measurements detected minimal stress-associated changes in the gut microbiome.

### Metaproteomic analysis reveals disrupted stress responses across diverse taxa in the male gut microbiome

Since conventional methods only revealed limited changes to the gut microbiome because of host stress, hidden changes in microbial function were investigated through a metaproteomic analysis of intestinal contents from GH and SI mice using DDA LC-MS/MS (S7; Fig S3A-D). In males, 14,408 microbial proteins were detected. Filtering and statistical analysis identified 811 significantly decreased and 319 significantly increased proteins in the SI group compared to GH controls (*q*<0.05, *abs(log_2_(fc))*>2; S7; Fig. S3E). Of the significantly altered proteins, 215 proteins were detected only in the GH mice and 54 were detected only in SI mice (S7).

Although 16S rRNA gene analysis revealed limited stress-associated changes in microbial composition, metaproteomics identified 73 significantly altered bacterial stress-response proteins in male SI mice (Fig. S3F). These proteins were assigned to diverse taxa, particularly within the Bacillota phylum, and included proteins involved in chaperone activity, protein quality control, oxidative stress response, and redox regulation (Fig. 3A & S3F). Notably, SI-induced host stress appears to remodel bacterial stress response proteins in specific pathways, and not through a general activation of all stress response mechanisms: SI stress was associated with an increased abundance of chaperone proteins *GroEL* and *ClpB*, and well as subunits of the Clp protease complex. In contrast, other molecular chaperones, including *HtpG* and *DnaK*, were decreased, along with multiple reactive species scavenging proteins, particularly rubrerythrin family proteins. This pattern is also observed within individual taxa. *Petralouisia muris*, *Anaerocolumna sedimenticola*, and *Anaerostipes rhamnosivorans* each demonstrated an increased abundance of chaperone ClpB or chaperone GroEL proteins together with decreased expression of oxidative stress response proteins (Fig. 3B-D).

**Figure 3.**
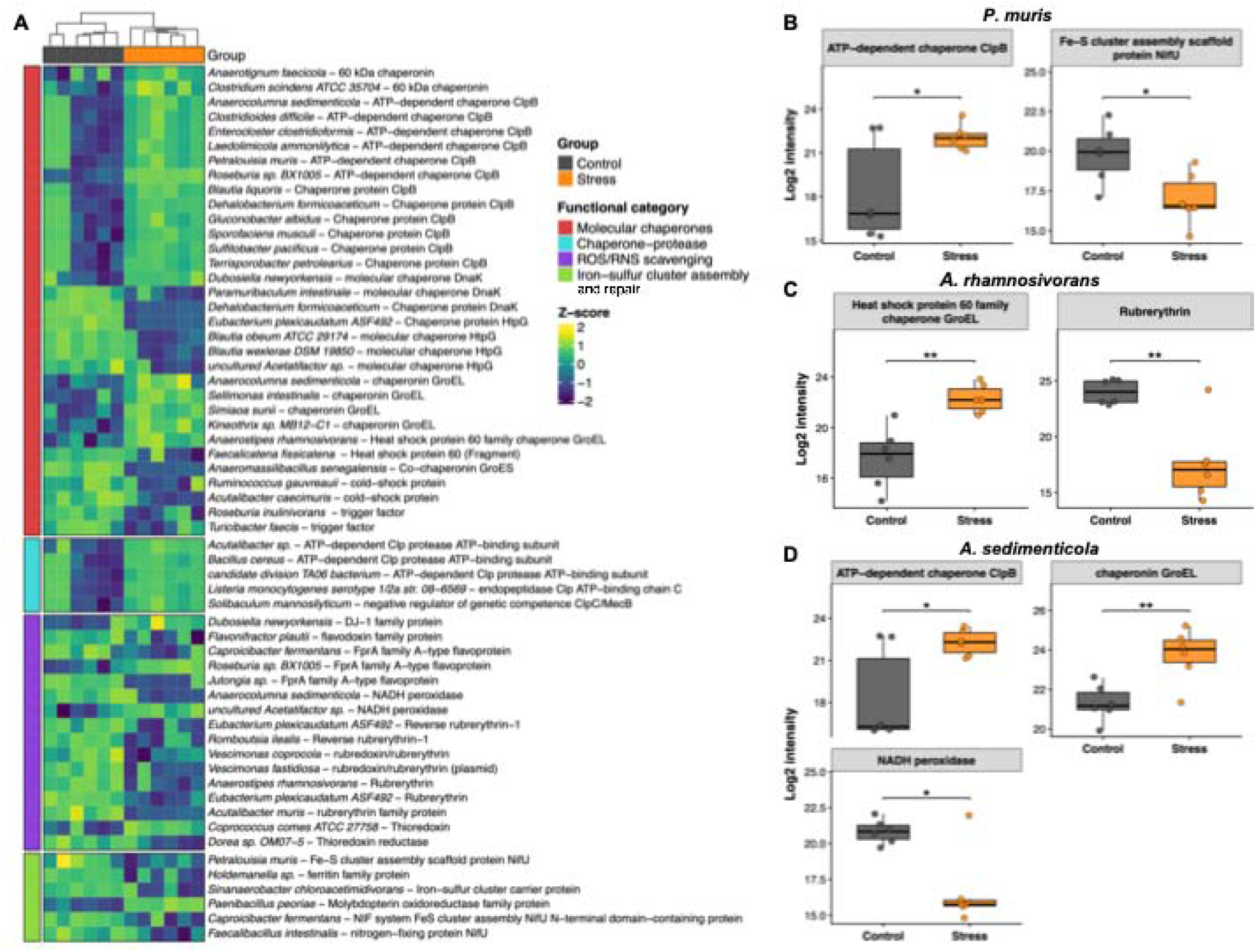
Metaproteomics reveals disrupted stress responses across diverse taxa in the male gut microbiome. **A** Significantly altered bacterial stress proteins (n=6 per group) identified using a metaproteomic analysis. Color-coded stress response functional categories are shown one the left. Z-score normalized intensities shown in purple (negative) to yellow (positive). **B-D** Box plots demonstrating the significant selective increase in certain chaperone proteins and decrease in general redox proteins across 3 taxa. Statistical significance was calculated using a weighted linear model and empirical-Bayes moderated t-test method through the limma R package (version 3.66.0). Raw *p*-values were adjusted using Benjamini-Hochberg (BH) false discovery rate (FDR) adjustment (*q*-value). Significance was defined as *q*<0.05 and an absolute Log2(fold change) ≥ 2: \**q*<0.05, \*\**q*<0.01. Whiskers extend to the most extreme values within 1.5X the interquartile range.

### Social isolation stress results in a specialized metabolic expansion of *Petralouisia muris* in the female gut microbiome

Metaproteomic analysis was also performed on intestinal contents from female GH and SI mice using DDA LC-MS/MS (S8; Fig. S4A-D). Across female large intestine samples, 11,414 proteins were detected. Filtering and statistical analysis revealed 371 significantly decreased and 232 significantly increased proteins in SI females compared to GH controls (*q*<0.05, *abs(log_2_(fc))*>2; S8; Fig. S4E). Among significantly altered proteins, 165 proteins were detected only in GH mice and 135 were detected only in SI mice (S8).

Within the significantly altered female metaproteome, 12 genera accounted for 53% of significant protein identifications (Fig. S4F). In this dataset, the genus *Petralouisia* was represented by a single species, *P. muris*, which had the highest number of significantly altered proteins among detected taxa, with 52 significant proteins (Fig. 4B). Isolated from the murine gut, *Petralouisia muris* is the type strain for the *Petralouisia* genus. Of the 52 significant *P. muris* proteins, 49 proteins were significantly more abundant in SI mice than in GH controls (Fig. 4A). These stress-enriched proteins spanned a variety of metabolic and cell cycle functions, suggesting a functional expansion of this taxon in response to host stress (Fig. 4B).

**Figure 4.**
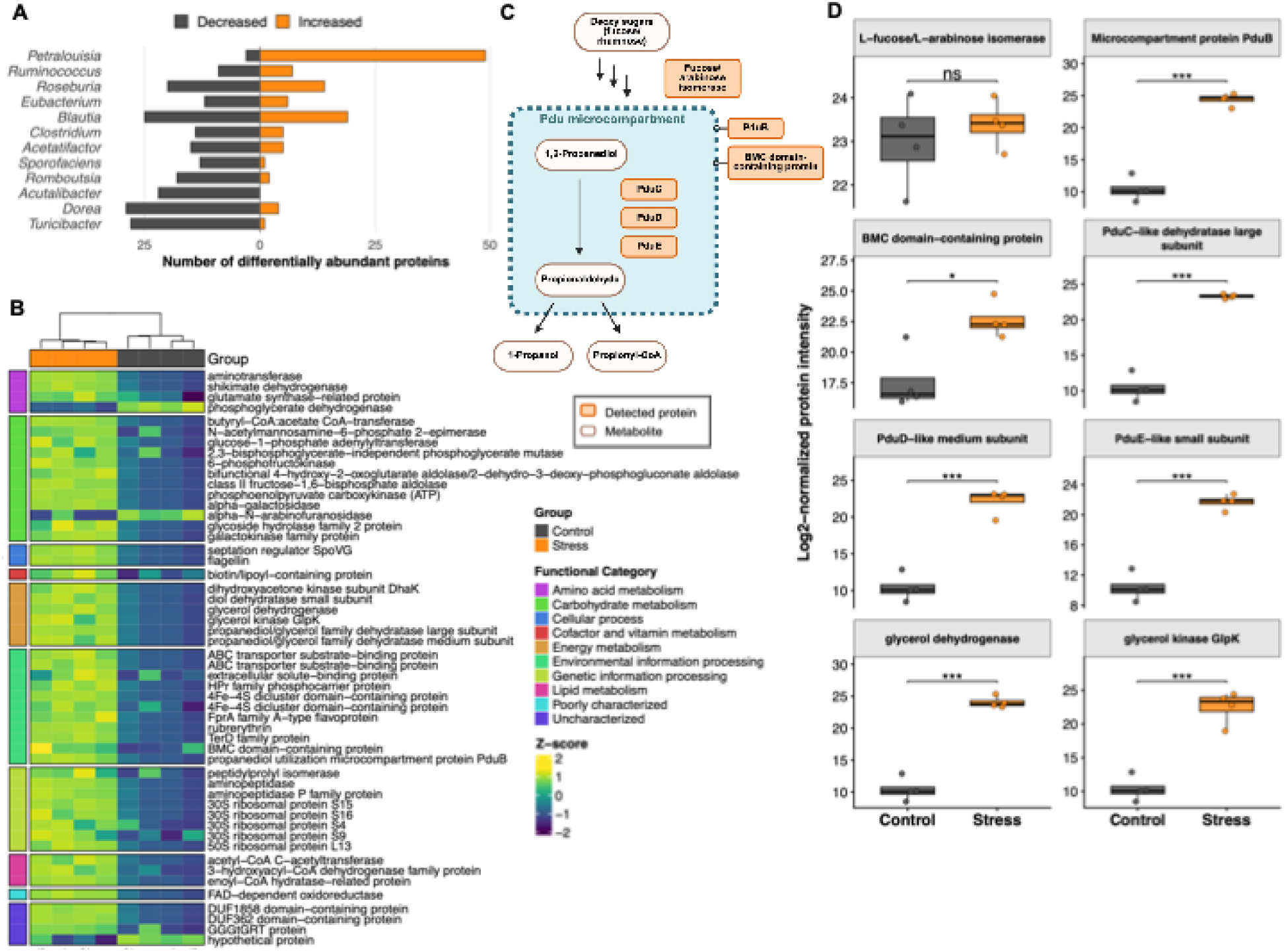
Stress results in a specialized expansion of *Petralouisia muris* in the female gut. **A** Horizontal bar chart showing the distribution of significantly altered proteins in the top 12 most abundant genera by protein count. **B** Significantly altered proteins from *P. muris* (n=4 per group) and their metabolic groupings. Color-coded functional categories are shown one the left. Z-score normalized intensities shown in purple (negative) to yellow (positive). **C** Simplified steps of the 1,2-propanedoil degradation pathway. Proteins detected in the *P. muris* dataset are highlighted in orange. Metabolites in the pathway are highlighted in tan. **D** Detected *P. muris* proteins involved in 1,2-propanediol degradation. Statistical significance was calculated using a weighted linear model and empirical-Bayes moderated t-test method through the limma R package (version 3.66.0). Raw *p*-values were adjusted using Benjamini-Hochberg (BH) false discovery rate (FDR) adjustment (*q*-value). Significance was defined as *q*<0.05 and an absolute Log2(fold change) ≥ 2: not significant (ns), \**q*<0.05, \*\**q*<0.01, \*\*\**q*<0.001. Whiskers extend to the most extreme values within 1.5X the interquartile range.

A member of the *Lachnospiraceae* family, *P. muris* is one of the 20% of all bacterial species that produce and use bacterial microcompartments (BMCs) for the compartmentalization of specialized metabolic functions, such as propanediol utilization (Pdu), to gain a competitive advantage for specialized niches in the gut (Fig. 4C)^51^. Within the detected *P. muris* proteome, 6 proteins directly associated with 1,2-propanediol utilization and 2 proteins associated with the closely related glycerol metabolism pathway were identified (Fig. 4D). Seven of these 8 proteins were significantly enriched in SI mice (*q*<0.05, *abs(log_2_(fc))*>2; Fig. 4D). The single protein that was not significantly enriched between conditions was an L-fucose/arabinose isomerase that is likely positioned upstream in the Pdu pathway and contributes to the degradation of deoxy sugars into 1,2-propanediol (Fig. 4C). Among the stress-enriched proteins, 2 were major structural components of the BMC: microcompartment protein PduB and the BMC domain-containing protein. Three additional stress-enriched proteins were components of the multienzyme complex propanediol dehydrogenase (PduCDE), which catalyzes the conversion of 1,2-propanediol to toxic propionaldehyde, a key step within the Pdu pathway. The remaining 2 stress-enriched proteins were assigned to the glycerol utilization pathway, which is also suggested to be Pdu-mediated^52,53^.

### Social isolation reshapes arabinoxylan and glucuronide metabolic capacity

Among the significantly different proteins between the SI-stress and GH control groups, 32 proteins in males and 26 proteins in females were assigned glycoside hydrolase activity (Fig. S5A & B). To further define the predicted carbohydrate-active functions of these proteins, predicted glycoside hydrolases were annotated with dbCAN3^54^, assigned to glycoside hydrolase families using the CAZy database^55^, and mapped to enzyme commission (EC) numbers^56^ (Fig. S5A & B; S7 & S8).

In males, 11 of the differentially abundant glycoside hydrolases were predicted to have β-xylosidase or α-L-arabinofuranosidase activity (EC 3.2.1.37/3.2.1.55; Fig. 5A). Of these, 8 were significantly decreased in SI mice compared to GH controls. These enzymes release xylose and arabinose monomers from arabinoxylan and related polysaccharides. Following liberation from the parent polysaccharide, xylose and arabinose can undergo further microbial metabolism via xylose isomerase and L-arabinose isomerase, respectively, and be assimilated into energy-producing metabolic pathways (Fig. 5B). In SI males, the expression of several L-arabinose isomerases and xylose isomerases were significantly decreased (*q*<0.05; S7; Fig. 5C & D). Together, the decreased abundances of both polysaccharide-degrading enzymes and downstream sugar isomerases indicate a reduced arabinoxylan-associated metabolic capacity in the SI male gut microbiome.

**Figure 5.**
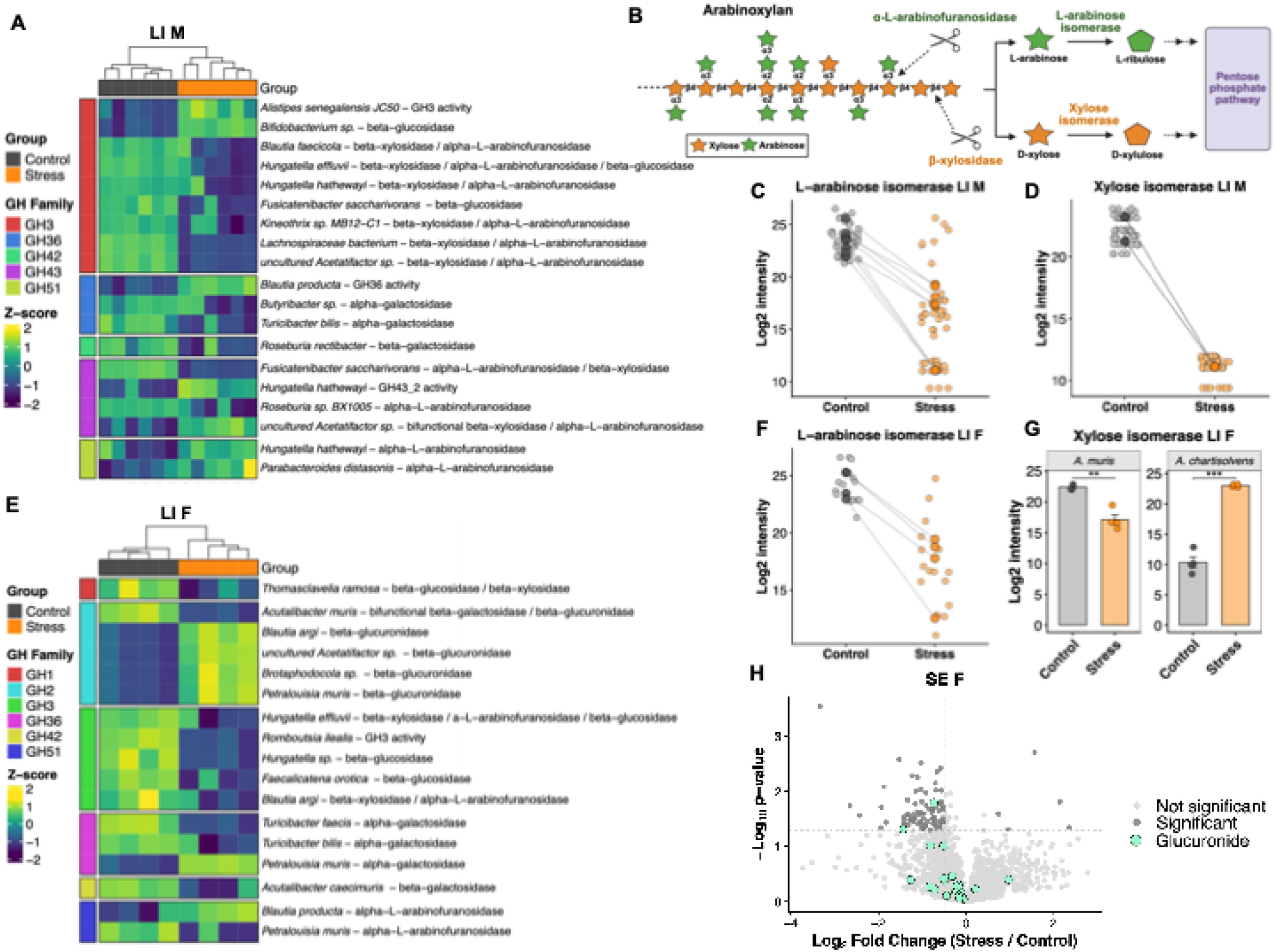
Social isolation reshapes arabinoxylan and glucuronide metabolism. **A** Selected significantly altered proteins in male mice microbiomes (n=6 per group) with predicted glycoside hydrolase activity. Color-coded glycoside hydrolase family groupings shown on the left. Z-score normalized intensities shown in purple (negative) to yellow (positive). **B** Simplified degradation pathway of arabinoxylan to precursors for the pentose phosphate pathway and microbial energy production. Enzymes detected in metaproteomic data here highlighted in green (arabinose-related) and orange (xylose-related). **C** Significantly decreased L-arabinose isomerase proteins in the male SI metaproteome. SI group replicates shown in light orange, mean SI expression values for each protein shown in dark orange, GH group replicates shown in light gray, mean GH expression values for each protein shown in dark gray. **D** Significantly decreased L-fucose isomerase proteins in the male SI metaproteome. **E** Selected significantly altered proteins in female mice microbiomes (n=4 per group) with predicted glycoside hydrolase activity. Color-coded glycoside hydrolase family groupings shown on the left. Z-score intensities shown in purple (negative) to yellow (positive). **F** Significantly decreased L-arabinose isomerase proteins in the female SI metaproteome. **G** Bar charts showing significantly altered xylose isomerase proteins in the SI female metaproteome. Error bars represent standard error of the mean. **H** Changes to the serum metabolite profile in female SI mice compared to the GH group. Candidate glucuronides are highlighted in green. Statistical significance for protein data was calculated using a weighted linear model and empirical-Bayes moderated t-test method through the limma R package (version 3.66.0). Raw *p*-values were adjusted using Benjamini-Hochberg (BH) false discovery rate (FDR) adjustment (*q*-value). Significance was defined as *q*<0.05 and an absolute Log2(fold change) ≥ 2: \**q*<0.05, \*\**q*<0.01, \*\*\**q*<0.001. Statistical significance of metabolites was calculated using Welch’s t-tests where significance was defined as *p*<0.05 and an absolute Log2(fold change) ≥ 0.5.

A similar pattern was observed in SI female mice. Five differentially expressed glycoside hydrolases were predicted to have β-xylosidase or α-L-arabinofuranosidase activity (EC 3.2.1.37/3.2.1.55; Fig. 5E). Four of these 5 proteins were significantly decreased in SI mice compared to GH controls. Proteins assigned L-arabinose isomerase activity were also significantly decreased in SI females (*q*<0.05; S8; Fig. 5F). In contrast, changes in expression of xylose isomerase proteins were taxon-specific: the abundance of xylose isomerase in *Acutalibacter muris* was decreased, whereas the abundance of xylose isomerase was significantly increased in *Anerobacterium chartisolvens* (*q*<0.05; S8; Fig. 5G).

In addition to changes in arabinoxylan-associated enzymes, SI female mice also showed increased abundance of proteins assigned β-glucuronidase activity across several Lachnospiraceae taxa (EC 3.2.1.23; Fig. 5E).

To determine whether SI was associated with changes in circulating glucuronide metabolites untargeted metabolomics was performed on serum from female mice (S9; Fig. 6H). Peak picking and compound annotation using Compound Discoverer software (Version 3.3, Thermo Fisher Scientific, Waltham, MA, USA) identified 1,989 metabolite features across the female serum metabolome. Among these features, 24 were annotated as candidate glucuronides (Fig. 6H). Of the circulating candidate glucuronides, 22 had lower abundance in SI mice than GH controls, although many did not reach statistical significance (*p*<0.05; S9; Fig. 6H). This emerging trend in circulating candidate glucuronide feature suppression combined with an increase of mGUS protein abundance in the female large intestine generates a mechanistic hypothesis for a potential stress-associated shift in glucuronide-related metabolism with prospective relevance to both dietary polysaccharide degradation and host-derived glucuronide processing.

**Figure 6.**
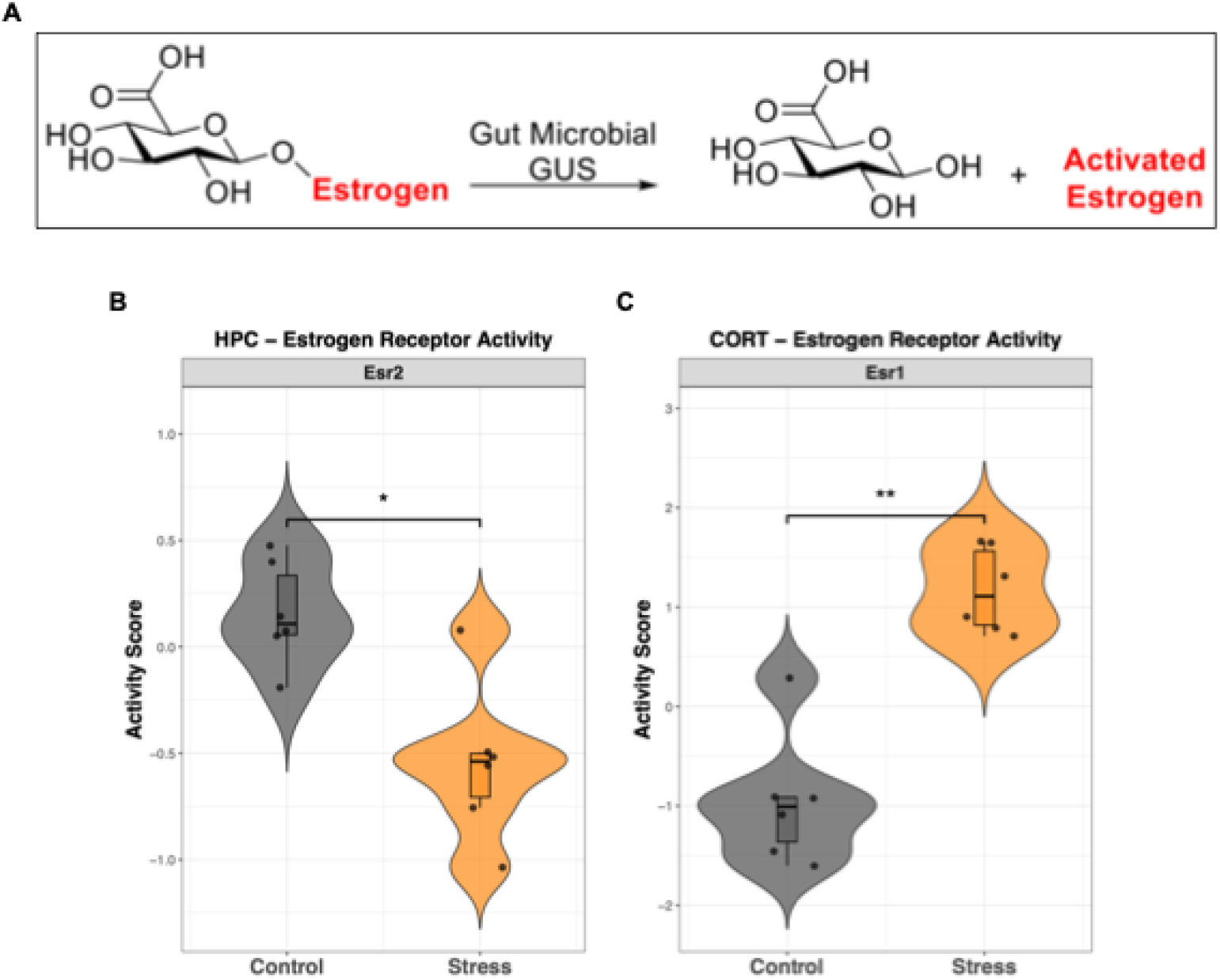
Isolated female mice demonstrate disruptions in estrogen response and signaling in the brain. **A** Predicted mechanism by which mGUS enzymes reactivate glucuronidated estrogen compounds. **B & C** Significant differences of estrogen receptor activity scores between SI and GH mice (n=6, per group) in the HPC and CORT brain regions. Transcription factor (TF) activity scores were predicted from protein targets with a multivariate linear model using the decoupleR R package (version 2.16.0). Statistical analysis of activity scores between experimental groups was conducted using a two-sided, unpaired Welch’s t-test. Raw *p*-values were adjusted using Benjamini-Hochberg (BH) false discovery rate adjustment (*q*-value). Significance was defined as *q*<0.05 and an absolute delta score ≥ 0.5: \**q*<0.05, \*\**q*<0.01. Whiskers extend to the most extreme values within 1.5X the interquartile range.

### Isolated female mice demonstrate region-specific disruptions in estrogen response and signaling in the brain

Microbial glucuronidases can act on glucuronidated host metabolites, including hormones, neurotransmitters, and metabolic byproducts, thereby re-activating aglycones that may then re-enter host signaling pathways^57^ (Fig. 6A). Among the candidate glucuronides detected in female serum, an estrogen metabolite candidate, estrone glucuronide, showed lower abundance in SI females compared to GH controls, although this change did not reach statistical significance (ns; S9; Fig. S6A). While not significant as an individual metabolite feature, the decrease in estrone glucuronide was consistent with the broader downward trend observed across circulating candidate glucuronides.

To evaluate whether SI-associated enrichment of mGUS expression coincided with changes in host signaling pathways, transcription factor (TF) activities were estimated from female hippocampal and cortical proteomic profiles using a Discriminant Regulon Expression Analysis^58^ (S2& S3). In the hippocampus, 49 TFs were detected, including 13 that were significantly decreased and 13 that were significantly increased in SI mice compared to GH controls (*q*<0.05; S2; Fig. S6B). In the cortex, 18 TFs were detected, including 3 that were significantly decreased and 7 that were significantly increased in SI compared to GH mice (*q*<0.05; S3; Fig. S6C). Among ligand-regulated TFs, inferred Estrogen Receptor 2 (*Esr2*) activity was significantly decreased in the hippocampus of SI mice, whereas inferred Estrogen Receptor 1 (*Esr1*) activity was significantly increased in the cortex of SI mice compared to GH controls (Fig. 6B & C).

To further asses SI-associated changes to estrogen signaling in the female brain, detected hippocampal and cortical proteins were annotated using the KEGG database^59^. In the hippocampus, 15 detected proteins mapped to the murine estrogen signaling pathway (KEGG: mmu04915; S2 & S3). Of these, the expressions of 8 hippocampal proteins were significantly altered between SI and GH mice, including 5 increased and 3 decreased proteins (Fig. S6D). In the cortex, 18 proteins mapped to the estrogen signaling pathway, including 4 proteins that were significantly enriched in SI mice (Fig. S6E). Many of the detected hippocampal and cortical proteins mapped to the KEGG estrogen signaling pathway are sensitive to psychological stress and play an overlapping role in other stress response pathways^60^. However, the abundance of proteins coupled with activated estrogen receptor complexes (*Gnai2*)^61^ and directly downstream of estrogen receptor activation (*Ctsd*)^62^ were significantly altered; further supporting a disruption in estrogen response (*q*<0.05, *abs(log_2_(fc))*>2). Together, these findings present a biologically relevant stress-associated gut microbial enzymatic signature that coincides with region-specific changes in brain estrogen receptor activity and signaling in female mice.

## Discussion

Although the GMBA has long been recognized as an important mediator of host stress responses, much of the field still relies on sequencing-based and inference-driven approaches to characterize the microbiome’s involvement in PS. In this study, we performed an integrated multi-omic analysis to investigate the effects of chronic SI stress on the GMBA of male and female mice. Despite the absence of an overt behavioral change, SI produced host molecular and microbial signatures that are consistent with a biologically meaningful stress response and patterns observed in disorders and neurological diseases. Conventional approaches, including 16S rRNA gene sequencing and targeted SCFA analysis, revealed minimal effects and did not capture the extent of stress-associated remodeling. In contrast, metaproteomics uncovered changes in microbial stress-responses, bacterial ecology, and polysaccharide-degrading capacity. In females, these functional shifts were accompanied by emerging serum metabolomic and brain proteomic patterns suggestive of potential downstream consequences for host estrogen signaling. Together, these findings demonstrate that stress can functionally remodel the GMBA even without a robust behavioral phenotype, highlighting the importance of function-based analyses for detecting subtle host-microbiome interactions.

Unlike stress models that rely on physical or unpredictable external pressures, SI models chronic and etiologically relevant PS^26^. However, the behavioral effects of SI depend on experimental variables including mouse age, strain, sex, and treatment duration^63–68^. Here, adult male and female C57BL/6J mice did not exhibit significant behavioral changes following 6 weeks of SI. Behavioral characterization was limited to the forced swim test and weekly body weight measurements. While these provided insight into depressive-like behavior and systemic effects of SI on diet and feeding, we did not include a behavioral assay designed specifically to assess anxiety-like behaviors. Therefore, the SI paradigm did not induce a severe depressive phenotype or systemic physiological change, although other behavioral consequences cannot be excluded. Importantly, the absence of robust behavioral changes does not imply the absence of biological stress. Proteomic analyses of the hippocampus and cortex revealed molecular patterns consistent with known stress response signatures. This distinction is important because many individuals experience chronic stress without meeting behavioral or diagnostic thresholds for stress-related disorders. Therefore, this SI paradigm provides a valuable model for identifying early or subtle molecular alterations that may precede more severe behavioral or physiological dysfunction.

A large body of evidence links PS to altered gut community composition, depletion of beneficial taxa, expansion of stress-tolerant or potentially pathogenic organisms, and changes in microbial metabolites such as SCFAs^18,20,69,70^. Based on this literature, we expected SI to produce detectable changes in bacterial composition and SCFA availability. However, 16S rRNA gene sequencing revealed no taxonomic shifts in females and limited changes in the males, while targeted SCFA analysis showed no significant SI-associated differences. Potentially, pooling cecal and colon microbial contents may have contributed to these limited effects. The intestinal tract contains distinct region-specific bacterial communities^71^, and subtle stress-associated changes within either the cecum of colon may have become obscured by combining them. Still, these findings emphasize that this SI stress paradigm produced minimal robust changes in community composition of SCFA concentrations, and that these analyses were insufficient to reveal the full extent of the stress-associated effects on the GMBA.

In contrast, metaproteomics provided complementary functional resolution, revealing stress-associated changes within the microbiome. In males, SI was associated with increased abundance of bacterial chaperone and heat shock proteins, including ClpB and GroEL, coupled with a broad decrease in redox proteins. Given the established link between PS and immune signaling in the gut^15^, the reduction in general redox proteins was unexpected. However, chaperone proteins also play a central role in bacterial survival in challenging environmental conditions, such as the inflamed host intestine^72^. These results suggest that the male metaproteome was affected by SI, even though taxonomy and SCFA concentrations remained largely unchanged. This suggests that changes in bacterial stress protein abundances were likely due to host stress signaling in the intestinal lumen, and nutrient availability pressures. In females, metaproteomics revealed a stress-associated expansion of *Petralouisia muris* proteins, including proteins involved in BMC-mediated propanediol utilization. BMCs compartmentalize specialized metabolic reactions and can confer competitive advantages during specific nutrient availability or inflammatory conditions, and studies have shown that enteric pathogens can outcompete commensal taxa through BMC-associated pathways in the inflamed intestine^51,53,73^. Enrichment of this pathway therefore suggests that SI stress may alter ecological opportunities within the gut, potentially benefitting bacteria capable of metabolizing propanediol or glycerol. It is tempting to speculate that with longer isolation, or in a more stress-susceptible cohort, pathogenic bacteria might similarly exploit BMC-associated metabolisms, contributing to dysbiosis and disease progression. Overall, while 16S rRNA gene sequencing and SCFA analysis did not capture the complete picture of stress-associated microbiome disruption, metaproteomics revealed that SI altered microbial physiology and ecological strategy in sex-specific ways.

One primary mechanism through which the gut microbiome interacts with the host is the degradation of dietary polysaccharides and host-associated glycans^74,75^. In homeostatic systems, gut bacteria metabolize dietary fibers and host glycans into compounds that support both microbial growth and host cell metabolism^74,75^. Subsequently, the carbohydrate-active enzyme (CAZyme) repertoire of the gut microbiome is increasingly recognized as a functional signature of host-microbiome health, and disruption in CAZyme profiles have been implicated in neurological and stress disorders^76–78^. Here, SI-associated stress altered the abundance of multiple arabinoxylan-degrading glycoside hydrolases in a sex-specific manner. In males, SI was associated with decreased abundance of glycoside hydrolase family 3 arabinoxylan-degrading enzymes, increased abundance of family 51 arabinoxylan-degrading enzymes, and mixed changes among family 43 enzymes. In females, SI was associated with decreased abundance of glycoside hydrolase families 1 and 3 arabinoxylan-degrading enzymes, as well as mixed effects on family 51 enzymes. These findings indicate that SI-induced PS does not uniformly suppress or enhance arabinoxylan degradation potential across taxa but instead reshapes metabolic capacity of the microbiome as a whole.

These results align with evidence that neurological and stress-related disorders are associated with altered microbial carbohydrate metabolism^77,79^. For example, a recent study on major depressive disorder (MDD) reported increased expression of plant-associated CAZymes in patients with MDD compared to healthy controls, including enrichment of glycoside hydrolase family 51 and glycoside hydrolase family 43 subfamilies as predictive CAZyme markers of MDD^77^. Conversely, a study of Parkinson’s disease (PD), a neurological disorder in which chronic stress is a factor in disease onset and progression^80^, reported broadly decreased glycoside hydrolase activity across global PD cohorts compared to healthy controls^78^. Together, these findings indicate that the SI paradigm disrupted the CAZyme profile in a manner consistent with clinical neurological and stress-related disorders, even without an overt behavioral stress phenotype.

To our knowledge, stress-associated alterations in mGUS activity have not been reported, even though these enzymes are increasingly recognized as important mediators of host-microbiome metabolic interactions^57,81^. In the gut, mGUS enzymes deconjugate host-derived glucuronides that enter the intestine through biliary excretion, reactivating aglycones that can be reabsorbed and returned to circulation^82^. Through this process, mGUS enzymes can influence the systemic availability of host-derived compounds, including neurotransmitters and hormones^57,81^. Here, SI females showed increased mGUS abundance in the gut microbiome alongside a decreasing trend in circulating glucuronide candidate features. While not all glucuronide features reached statistical significance, the overall pattern was consistent with an increase in glucuronidase activity.

These findings are particularly notable because they identify a previously undescribed stress-associated increase in mGUS abundance coinciding with altered estrogen receptor activity in the female brain. Specifically, inferred Estrogen Receptor 2 (*Esr2*) activity decreased in the hippocampus, whereases inferred Estrogen Receptor 1 (*Esr1*) activity increased in the cortex. Previous work has shown that microbial reactivation of glucuronidated endobiotics can influence host physiology. For example, gut microbial reactivation of estrogen metabolites has been implicated in breast and ovarian cancer^83^, and Simpson *et al*. found that mGUS modulates serotonin availability, with well-known implications for depression^57^. The observed estrogen receptor pattern is also broadly consistent with evidence that chronic stress can selectively suppress brain *Esr2* expression and increase *Esr1* expression in rodent models^84^. Increased circulating estrogen metabolites have also been associated with decreased *Esr2* expression^85^. Together, these results support a novel mechanistic hypothesis in which stress-associated microbial metabolism disrupts female neuroendocrine signaling in a manner consistent with known physiology of stress disorders and neurological diseases. This is especially notable because the SI paradigm modeled a subclinical behavioral response. Although this study did not establish a causal pathway through which host stress alters microbial metabolism, the convergence of metaproteomic, serum metabolomic, and host proteomic evidence provides a strong framework for determining how host stress signaling interfaces with the microbiome in SI models.

To our knowledge, this is the first study to characterize the GMBA in a chronic stress model with subclinical behavioral outcomes. Overall, these findings support a model in which chronic SI produces biologically meaningful functional changes to the GMBA apart from overt behavioral changes. However, a complete picture of SI-induced GMBA remodeling was not detectable through conventional sequencing-based analyses or SCFA quantification. Instead, disruptions in microbiome-host interactions emerged through functional multi-omic profiling. Specifically, we identified increased mGUS abundance as a novel microbial response to PS, with potential consequences for estrogen signaling in the female brain. These findings reveal a notable parallel between the microbiome and the brain during chronic stress. Just as molecular and cellular adaptations in stress-sensitive brain regions occur before detectable behavioral changes^86^, microbial communities can undergo functional remodeling despite maintaining a relatively stable taxonomic composition. This work underscores the need to move beyond inference-based analyses of GMBA interactions and suggests that functional microbiome profiling may be especially important for understanding PS physiology in individuals experiencing chronic stress without a diagnosable behavioral or clinical phenotype.

## Methods

### Experimental design

#### Ethics declaration

The animal study was approved by the Institutional Animal Care and Use Committee at Baylor University (Protocol No. 2025019-2).

#### Animals and social isolation

Following a 4-week acclimation period, male mice (n=12; 8 weeks old) and female mice (n=12; 8 weeks old) were randomized into 2 groups of 6 animals, per sex, for a total of 4 groups of mice. For male (n=6) and female (n=6) control groups, mice were housed in groups of 3 mice of matched sex per cage. Social isolation mice (n=6 male and n=6 female) were housed individually. The treatment period lasted 6 weeks and all mice had *ad libitum* access to sterilized food and an automatic central water system. All mice were housed at roughly 20 °C and 50% humidity, with a 12 h light-dark cycle, high-efficiency particulate air filters, and sterilized individually ventilated cages, wire food trays, bedding, and one cotton square per cage. To reduce visual stimulation and interaction, social isolation cages were wrapped in white masking tape. Physical health status was determined by observation for distress symptoms including hunched posture, changes in grooming habits, changes in breathing or mobility, and body weight. All mice were monitored 3 times a week for health status. Body weight was monitored weekly. No animals showed a decline in physical health through the duration of the study. All mice were handled regularly by the same researcher, 3 times a week during physical health monitoring and fecal sample collection, and once a week for body weight measurement.

#### Behavioral analysis

At the end of the 6^th^ treatment week, despair-like behavior detection was conducted using a 6-min forced swim test (FST; n=12 male, n=11 female)^87^. The first 2 min were a pre-test, and data were only collected during the final 4 min of the test. The total immobility time within the 4-min test was analyzed by 3 researchers blinded to the experimental group of each mouse. FST was conducted in the early morning while the mice were in their active period. Mice were forced to swim, once, for 6 min in a polypropylene 4 L beaker (Thermo Fisher Scientific, Waltham, MA, USA) filled with 3 L of water at 25 °C. The swim beaker was emptied and cleaned between each mouse. After the test, each mouse was placed in a clean cage, maintained at 37 °C, with blotting paper and monitored. The paper was changed, and the cage was cleaned after each mouse.

Significant differences between group immobility times were determined using a two-sample t-test where significance was defined as *p*<0.05. Twenty-four hours after FST all mice were euthanized. Serum, intestinal contents, colon tissues, hippocampi, and prefrontal cortexes were collected. For further information about animal husbandry, behavioral analysis, and euthanasia, please see supplementary materials.

#### Euthanasia and sample collection

Twenty-four hours after FST, all mice were euthanized by cervical dislocation. Cardiac puncture was performed directly post-mortem for blood sample collection. Blood was centrifuged 1,500 x *g*, 4 °C for 20 min. Serum was transferred to a clean tube, flash frozen, and stored at −80 °C. Large intestinal contents (pooled cecum and colon) were collected, flash frozen, and stored at −80 °C. Colon tissues were rinsed with a cold 1.2% NaCl solution, Invitrogen™ RNAlater™ Stabilization Solution (700 μL; Thermo Fisher Scientific, Waltham, MA, USA) was added, samples were flash frozen, and stored at −80 °C. Brains extracted immediately after euthanasia and rinsed with a cold 1.2% NaCl solution. The prefrontal cortexes and the hippocampi were immediately dissected on ice, flash frozen, and stored at −80 °C for subsequent analyses.

### Host proteomics

#### Preparation of brain samples

Approximately 15 mg of hippocampus and cortex tissue was used from each mouse (n=24). Tissues were minced using dissection scissors and further homogenized on ice with a Dounce homogenizer (A and B pestles 15 times, each; DWK Life Sciences, Vineland, NJ, USA) with cold 250 mM sucrose 1XPBS (200 μL). Homogenate was centrifuged (10,000 x *g*; 4 °C; 25 min). Supernatant protein concentrations were determined using a Pierce™ Bicinchoninic Acid (BCA) assay (Thermo Fisher Scientific, Waltham, MA, USA) read by a BioTek Cytation 5 Cell Imaging Multimode Reader (Agilent Technologies, Inc., Santa Clara, CA, USA). All samples were normalized to 1 mg/mL protein concentration.

Neat urea (75 mg) was added to the normalized tissue lysates (100 μL) in 1.5 mL tubes, followed by reduction using dithiothreitol (DTT; 1 M; 3 μL). Samples were incubated (50 °C; 45 min; 1200 rpm). After cooling to room temperature, iodoacetamide (IAA; 200 mM; 10 μL) was added and samples were incubated in darkness (45 min; 25 °C; 1200 rpm). After incubation, DTT solution (500 mM; 3 μL) was added and samples incubated (30 min; 30 °C; 1200 rpm). Next, ammonium bicarbonate (50 mM; 320 μL; pH 8) and CaCl_2_ (1M; 1 μL) were added and samples were digested for 16 h with Promega MS-grade trypsin (Thermo Fisher Scientific, Waltham, MA, USA) at a ratio of 1 μg trypsin for 50 μg sample protein (37 °C; 1200 rpm).

Following digestion, a C18 solid phase extraction was used for desalting. Briefly, Discovery® DSC-18 SPE Tubes (Sigma-Aldrich, Inc., St. Louis, MO, USA) were set up on a standard, 12-port Visiprep™ SPE Vacuum Manifold (Sigma-Aldrich, Inc., St. Louis, MO, USA). Under vacuum, the following was flowed through the columns: MeOH (3 mL), water with 0.1% trifluoroacetic acid (TFA; 2 mL), samples, and 95:5 water with 0.1% TFA: acetonitrile (ACN; 4 mL). Cleaned samples were eluted by 80:20 ACN: water with 0.1% TFA (1 mL) into new 1.5 mL tubes. Fractions were dried in an Eppendorf Vacufuge (Eppendorf North America, Enfield, CT, USA) and peptides were stored at −20 °C. Samples were reconstituted in ammonium bicarbonate (25 mM; pH 8; 40 μL; 5 min; 37 °C; 1000 rpm). Samples were transferred to Beckman Coulter polycarbonate ultracentrifuge tubes and ultracentrifuged (100,000 x *g*; 20 min; 4 °C). Peptide concentrations were measured using a Pierce™ fluorometric peptide quantification assay (Thermo Fisher Scientific, Waltham, MA, USA) and normalized to 0.25 μg/μL, transferred to 0.2 mL amber MS vials (30 μL; Thermo Scientific™ SureSTART™ 0.2 mL TPX Screw Top Microvial with Glass Insert) and stored at −20 °C until analysis by LC-MS/MS.

#### LC-MS/MS proteomics analysis

Host tissue peptide samples were blocked by sex, individually randomized within blocks, and analyzed on a Thermo Scientific Orbitrap Fusion Tribrid Mass Spectrometer with a Vanquish Neo analytical-flow LC system (Thermo Fisher Scientific, Waltham, MA, USA). The column was a Thermo Scientific C18 HPLC column (15 cm length, 75 μm inner diameter, a 100 Å pore size, and a 2 μm particle size; Thermo Fisher Scientific, Waltham, MA, USA). The ion transfer tube temperature and spray voltage were set to 300 °C and 2.5 kV, respectively. The following run method was used for global proteomics: 8 μL containing 2 μg of peptides were injected into the analytical column at 0.3 μL/min. The mobile phases consisted of 0.1% formic acid in water (A) and 0.1% formic acid in 80:20 acetonitrile: water (B) with the following gradient profile (min, %B): 0, 2; 2, 2; 100, 25; 20, 40; 1, 95; 1, 95. After every sample block, the column was washed with the following wash method (min, %B): 0, 2; 1, 2; 0.1, 95; 1, 95; 0.1, 2; 1, 2; 0.1, 95; 3.7, 95. The data-dependent acquisition mode was used to automatically trigger the precursor scan and the MS/MS scans. The MS^1^ spectra were collected at a scan range of 350-1400 m/z, a resolution of 120,000, and automatic gain control (AGC) target of 8 × 10^5^. For MS^2^, data-dependent acquisition (DDA) was performed using a 3 s cycle time. The most abundant precursor ions above an intensity threshold of 1 × 10^4^ and within a window of 1.6 m/z were selected for fragmentation using quadrupole isolation and fragmented by higher-energy collisional dissociation (HCD) with a normalized collision energy (NCE) at 30%. The ion trap was used to collect MS^2^ spectra with a maximum AGC target of 1 × 10^5^, and a maximum ion injection time of 35 ms. Each parent ion was fragmented once before being dynamically excluded for 30 s using a parent ion tolerance window of 5 ppm.

#### Bioinformatic processing and statistical analysis

Raw (.raw) spectra files were processed and analyzed using an in-house bioinformatics pipeline. Briefly, raw files were converted to .mzML files using ThermoRawFileParser with the option of centroid smoothing^88^. The .mzML files were analyzed using MS-GF+ as a database search engine for identifying peptides using a standard target-decoy database search approach^89^. The FASTA file for the mouse protein database was downloaded from Uniprot (*Mus musculus*, accessed March 28, 2025). Two missed cleavages were allowed in fully and partially tryptic peptides. Peptide mass tolerance was set to 20 ppm, cystine alkylation was set as a static modification, and methionine oxidation was set as a dynamic modification. The identified peptide spectrum matches (PSMs) from the MSGF+ pipeline were rescored using an in-house rescoring algorithm—XPECTRA (not published). XPECTRA is an Extreme Gradient (XG) Boost-based classifier. Rescored PSMs were filtered based on a model-calculated *q*-value (xpec-q) < 0.05. The filtered PSMs were then rolled up to protein level using the Reference Rollup (RRollup) method^90^ implemented through a custom-built python script where an FDR of 5% and a minimum of 2 unique peptides were required for evidence.

To identify proteins with differential abundances between groups data processing for host hippocampus and prefrontal cortex occurred as follows. First, datasets were filtered to include only protein hits that were present in ≥75% of one group. Next, analysis continued in R where signal intensities were log_2_ transformed, and samples were median normalized. After normalization, missing values were imputed using a hybrid imputation strategy with the imputeLCMD R package (version 2.1)^91^. For any samples within a group that had missing values when ≥75% of that group’s samples contained the protein where determined “missing at random” and the missing samples in that group were imputed using the K-nearest neighbors (kNN) method. For any samples within a group that had missing values when <75% of that group’s samples contained the protein were determined “missing not at random” and the missing samples in that group were imputed using a minimum detection approach (MinDet). Finally, if all samples in a group had missing values for a protein, that protein was considered “absent” and the missing value in each sample for that protein was imputed with the lowest value (>0) detected across all proteins in that sample. After imputation, variance in weights was corrected and statistics were performed using a weighted linear model and empirical-Bayes moderated t-test method with the limma R package (version 3.66.0)^92^. P-values were adjusted (*q*-value) using the Benjamini-Hochberg (BH) false discovery rate (FDR) correction. Proteins with *q*<0.05 and an absolute log_2_ fold change of greater than 2 (abs(log_2_(fc))>2) were considered significant. In female brain samples, a discriminant regulon expression analysis (DoRothEA) was performed to estimate transcription factor activity with the decoupleR^58^ R package (version 2.16.0) using Multivariate Linear Model and a minimum number of 5 targets. Statistical significance of differentially active regulons between conditions was determined by a two-sided, unpaired Welch’s t-test, and *p*-values were adjusted using BH FDR correction. Significance was defined as *q*<0.05. Proteins belonging to the murine estrogen signaling pathway were annotated and grouped using Kyoto Encyclopedia of Genes and Genomes^59^ through the STRING database and web platform (version 12.0)^93^.

### Colon inflammatory markers

#### Preparation of colon samples

RNA was extracted from colon tissue (∼ 50 mg) from each mouse (n=24) using the E.Z.N.A ® Total RNA Kit I (Omega Bio-Tek Inc., Norcross, GA, USA) according to manufacturer’s instructions. Total RNA concentration was determined, and each sample was normalized to 5 ng/μL. Following normalization, RNA from each sample (50 ng) were pooled into 1 sample per group for a total of 4 samples. cDNA was synthesized from pooled samples with iScript Reverse Transcription Supermix (Bio-Rad Laboratories, Inc., Hercules, CA, USA). Within a 20 μL reaction volume, PCR reactions consisted of cDNA (30 ng), 1x SYBR Green (Thermo Fisher Scientific, Waltham, MA, USA), forward primer (0.4 μM), reverse primer (0.4 μM), and water (refer to SI4 for primer sequences).

#### qPCR analysis

Quantitative PCR was performed using a QuantStudio™ 6 (Thermo Fisher Scientific, Waltham, MA, USA) as follows: 1 cycle of 2 min at 50 °C then 95 °C; 40 cycles of sample denaturation at 95 °C for 15 sec, annealing at 55 °C for 15 sec, and extension at 72 °C for 1 min. Amplification specificity was verified using a melt curve as follows: continuous hold at 95 °C for 15 sec (1.6 °C /sec ramp), 60 °C for 1 min (1.6 °C /sec ramp), and 95 °C for 15 sec (0.15 °C /sec ramp). β-actin was used as the house-keeping gene control and all samples were run in quadruplicate.

#### Data processing and statistical analysis

Measured cycle threshold (Ct) values from each sample were normalized with Ct values from the house-keeping gene. Once all Ct values were normalized, a two-samples t-test and the Ct fold change (expressed as 2^-ΔΔCt) between sex-matched control mice and stressed mice was performed and determined. A significant difference in expression was defined by a *p*<0.05.

### Microbiota full-length 16S rRNA gene long-read sequencing

#### Preparation and sequencing of microbial DNA

Large intestinal gut content (∼15 mg) was used from each mouse (n=24). Total DNA was extracted from each sample using the ZymoBIOMICS™ DNA Miniprep Kit (Zymo Research, Irvine, CA, USA) and normalized to 1 ng/μL using the Qubit™ dsDNA Quantification Assay Kit (Thermo Fisher Scientific, Waltham, MA) and a Qubit™ 4 Fluorometer (Thermo Fisher Scientific, Waltham, MA). Samples were stored at −20 °C until library preparation. For library construction, purified DNA (2 ng) from each sample was processed using the *Quick*-16S Full-Length Library Prep Kit with Primer Set 1 (Zymo Research, Irvine, CA, USA) and amplified by qPCR on a QuantStudio™ 6 (Thermo Fisher Scientific, Waltham, MA, USA). Nuclease-free water was included as a negative control, and DNA from the mixed bacterial suspension provided by Zymo Research in the library prep kit was used as the positive control. Resulting amplicons were pooled into one library. Amplicon DNA from the library (200 fmol) was prepared for sequencing by the Ligation Sequencing Kit V14 (SQK-LSK114) (Oxford Nanopore Technologies). The purified DNA library (50 fmol) was sequenced by nanopore sequencing on a FLO-MIN114 (Oxford Nanopore Technologies, Oxford, UK) flow cell and MinION portable sequencer (Oxford Nanopore Technologies, Oxford, UK).

#### Bioinformatic processing and statistical analysis

Basecalling was deactivated during sequencing since sequencing was completed using custom barcodes and primers (Zymo long read 16S prep kit – D6450-PS1). After sequencing, reads in .pod5 raw files were basecalled using Dorado_dna_r10.4.1_e8.2_400bps_hac@v5.2.0 model (Oxford Nanopore Technologies basecaller; https://github.com/nanoporetech/dorado) to generate FASTQ files. The .fastq files were demultiplexed according to Zymo’s instructions via python script (accessed from https://github.com/calacademy-research/minibar/blob/master/minibar.py). Sequences were filtered based on a q-score threshold of ≥ 9. Using an in-house shell script, required minimum and maximum sequence lengths were 950 bp and 1750 bp, respectively.

To assign taxonomy and abundance estimations, filtered FASTQ files were processed with EMU (version 3.5.1)^94^, using an in-house custom EMU database curated from a 16S rRNA gene full length sequence databases including SILVA (138.2 release)^95^, GTDB (226 release)^96^, and MIMt (M2c_25_10 release)^97^. EMU output .tsv files containing the taxonomically assigned feature abundance table and corresponding metadata were imported into R, integrated using the phyloseq package (version 1.54.2) for downstream analyses.

Apha diversity metrics were calculated, and group differences were assessed using the non-parametric Wilcoxon rank-sum test where significant values were determined by *p*<0.05. Beta diversity was estimated by Bray-Curtis dissimilarity distances, and the overall community differences were evaluated using permutational multivariate analysis of variance (PERMANOVA; adonis2 function in R package vegan, version 2.7.5) with 9,999 permutations. A significant effect size was defined as *p*<0.05. Sample clustering and separation were then visualized using a principal coordinates analysis (PCoA) where percent variances of PC1 and PC2 are explained by x- and y-axes. Microbial composition of each group was summarized by relative abundance bar plots at the genus level generated by ggplot2 (version 4.0.3). Differential abundance analysis of genera between the two conditions was performed on the 16S rRNA gene-derived relative abundance data using MaAsLin2^50^ in R. Briefly, taxa with a minimum abundance of 0.0001 or prevalence <50% across samples were excluded prior to analysis. The remaining data were log-transformed and analyzed using linear models with Experimental Group as a fixed effect. P-values were adjusted using the BH FDR correction method so that significant differences were defined by *q*<0.05.

### Targeted SCFA metabolomics

#### Preparation of samples for SCFA quantification

Large intestinal content (50 mg) was used from each mouse (n=24) and homogenized in NaOH (0.5 M; 20 μL) and ice cold MeOH (480 μL) by vortexing. Samples were lysed by adding silca beads (∼200µL) to each sample and bead beating with an Omni Bead Ruptor Elite (Omni International, Inc., Kennesaw, GA, USA; 4.20 m/s; 45 s) immediately followed by a 5 min rest on ice, repeated for 3 lysing cycles. Lysate was centrifuged (17,500 x *g*; 30 min; 4 °C). Supernatants were transferred to new tubes. To precipitate protein, ice cold MeOH (80% v/v; 400 µL) was added, samples were vortexed for 30 sec and incubated (−20 °C; 30 min). Following incubation, proteins were pelleted by centrifugation (18,213 x *g*; 10 min; 4 °C). Supernatants (700 µL) were transferred to a new tube and dried using an Eppendorf Vacufuge (Eppendorf North America, Enfield, CT, USA). Dried samples were resuspended in NaOH (5 mM; 1 µL). Samples were transferred to glass vials and derivatization was performed using hexyl chloroformate. Briefly, a hexanol/pyridine solution (3:2 v/v; 500 µL) was added to each sample and mixed thoroughly. Hexyl chloroformate (100 µL) was added, and each sample was shaken, vented, and incubated (60 °C; 60 min). After derivatization, a hexane extraction (300 µL) was performed on each sample to isolate the derivatized compounds in the organic layer. A hexane blank was prepared. To enable quantification, a calibration curve was created for each acid by preparing acid standards in starting concentrations of 0 µg/mL, 5 µg/mL, 10 µg/mL, 25 µg/mL, 50 µg/mL, 100 µg/mL, 250 µg/mL, and 500 µg/mL.

#### GC-MS targeted analysis

Samples were randomized and analyzed using a Thermo Scientific ISQ 7000 Mass Spectrometer (Thermo Fisher Scientific, Waltham, MA, USA) with sample introduction accomplished via a Thermo Scientific AI 1310 autosampler (Thermo Fisher Scientific, Waltham, MA, USA). The column was a Thermo Fisher TG-5MS capillary column (Thermo Fisher Scientific, Waltham, MA, USA; 5% diphenyl/95% dimethyl polysiloxane with dimensions 30 m x 0.25 mm internal diameter and 0.25 µm film thickness). Helium was used as the carrier gas at the constant flow rate of 1 mL/min. Samples (0.5 μL) were injected using pressure-pulsed split conditions (10 psi) with a split ratio of 12:1. The column temperature was set to 50 °C for 2 min, increased to 70 °C by 10 °C/min, 85 °C by 3 °C/min, 110 °C by 5 °C/min, then to 290 °C by 30 °C/min and maintained for 5 min. The total run time was 25 min. Acquisition was performed in full-scan mode with a mass range of 30-300 m/z. The injector, transfer line, and ion source temperatures were used at 260 °C, 290 °C, and 230 °C, respectively. Data processing was performed using Chromeleon (version 7, Thermo Fisher Scientific, Waltham, MA, USA).

#### Data processing and statistical analysis

To account for decomposition of derivatized compounds over instrument run time, a decay correction was calculated and performed for each sample as described in S6. Statistical analysis was performed by using a two-sample t-test and statistical significance was determined by *p*<0.05.

### Metaproteomics

#### Preparation of large intestine samples

Large intestinal content (∼50 mg) was used from each mouse (n=12 male, n=8 female). Each sample was homogenized in cold 1X PBS (500 μL) by vortexing. Bacterial cells were lysed via bead beating as described above. Protein concentrations were determined by BCA assay and normalized to 1mg/mL. Global metaproteomic sample preparation on large intestinal content for LC-MS/MS was completed as described for host proteomic analysis with the following changes: dried peptides were resuspended in a 0.1% formic acid, 2% ACN and 95% H_2_O solution (v/v/v; 40 μL), and resuspended samples were centrifuged (18,213 x *g*; 30 min; 4 °C).

#### LC-MS/MS metaproteomics analysis

Samples were blocked and randomized as described for proteomic analysis of host tissues. Samples were run on a Thermo Scientific Orbitrap 480 Mass Spectrometer (Thermo Fisher Scientific, Waltham, MA, USA) with a Vanquish Neo analytical-flow LC system (Thermo Fisher Scientific, Waltham, MA, USA). The column was a Thermo Scientific C18 HPLC column (50 cm length, 75 μm inner diameter, a 100 Å pore size, and a 2 μm particle size; Thermo Fisher Scientific, Waltham, MA, USA). The ion transfer tube temperature and spray voltage were set to 280 °C and 2.5 kV, respectively. The following run method was used for global proteomics: 1 μg of peptides (4 μL of 0.25 μg/μL peptide concentration) was injected into the analytical column at 0.3 μL/min. The mobile phases consisted of 0.1% formic acid in water (A) and 0.1% formic acid in 80% acetonitrile/20% water (B) with the following gradient profile (min, %B): 0, 1; 0.5, 3; 120, 36; 0.5, 99; 9, 99. After every sample block, the column was washed with the following wash method (min, %B): 0, 1; 3, 1; 0.5, 99; 5, 99; 0.5, 1; 5, 1; 0.5, 99; 5, 99; 0.5, 1. The DDA mode was used to automatically trigger the precursor scan and the MS^2^ scans. The MS^1^ spectra were collected at a scan range of 375-1475 m/z, a resolution of 120,000, and AGC target of 3 × 10^6^. For MS^2^, DDA was performed using a 3-second cycle time. The most abundant precursor ions above an intensity threshold of 5 × 10^3^ and within a window of 1.6 m/z were selected for fragmentation using quadrupole isolation and fragmented by HCD with a NCE at 30%. The ion trap was used to collect MS^2^ spectra with a maximum ion injection time set to auto at a resolution of 15,000. Each parent ion was fragmented once before being dynamically excluded for 30 sec using a parent ion tolerance window of 10 ppm.

#### Bioinformatic processing and statistical analysis

Bioinformatic processing and analysis was completed as previously described for analysis of host tissues with the following changes: The database was curated from species-level 16S rRNA gene sequence data from this study. FASTA files for each species detected were downloaded from Uniprot and combined to form one FASTA file per experimental group by sex for a two-step target-decoy database search approach as described previously^98^. The static modification was cystine alkylation and the dynamic modification was oxidation of methionine. Peptides were filtered using an MSMS Score of 8 and a minimum of 2 peptides were required for protein level identification where at least 1 peptide was required to be unique to that protein. Downstream processing and statistical analyses were completed as described for host proteomes. For carbohydrate-active enzymes, functional annotation was performed using dbCAN3^54^, CAZy^55^, and ENZYME^56^ databases.

### Untargeted metabolomics

#### Preparation of serum samples

Serum from each female mouse (50 μL; n=10) was thawed on ice. Proteins were precipitated by adding ice cold methanol (150 μL), vortexing, and centrifugation (18,213 x *g*; 20 min; 4 °C). Following centrifugation, the supernatant (150 μL) was transferred to a new tube and dried at room temperature using a vacufuge as previously described. Samples were reconstituted in 95:5 water: methanol (v/v; 50 μL). Samples were immediately analyzed after reconstitution. Quality control (QC) samples consisted of a pool of all samples (3 μL each sample). Blanks consisted of 95:5 water: methanol (v/v).

#### LC-MS/MS metabolomics analysis

Samples were randomized and analyzed using a Thermo Exploris 120 with a Vanquish Neo analytical-flow LC system (Thermo Fisher Scientific, Waltham, MA, USA). The column was a Thermo Scientific Hypersil GOLD aQ C18 Polar Endcapped HPLC Column (15 cm length, 2.1 mm diameter, and a 1.9 μm particle size; Thermo Fisher Scientific, Waltham, MA, USA). The ion transfer tube temperature and spray voltage were set to 320 °C and static mode, respectively. The following run method was used for metabolomics: 2 μL of each sample (40 μL) was injected onto the analytical column at 300 nL/min. The mobile phases consisted of 0.1% formic acid in water (A) and 0.1% formic acid in methanol (B) with the following gradient profile (min, %B): 0, 0; 5, 50; 6, 98; 10, 98; 10.1, 0; 15, 0. Data were collected by DDA and run in negative ionization mode. MS^1^ spectra were collected at a resolution of 120,000 and the scan range was 70-1000 m/z over 15 min with an AGC target set to standard and a maximum injection time of 50 ms. For MS^2^ analysis, the top 4 most abundant precursor ions exceeding the intensity threshold of 2 × 10^4^ were selected for fragmentation using a quadrupole isolation window of 1.5 m/z. Fragmentation was performed by HCD with stepped normalized collision energies (NCE) of 30, 60, and 90%. MS^2^ spectra were acquired at a resolution of 30,000, AGC target set to standard, and a maximum injection time mode set to auto. Each parent ion was fragmented once before being dynamically excluded with auto mode.

#### Bioinformatic processing and statistical analysis

Feature detection, extraction, alignment, gap filling, and blank subtraction was performed using Compound Discoverer Version 3.3 (Thermo Fisher Scientific, Waltham, MA, USA). Features were annotated using the mzCloud spectral library, and mass lists Endogenous Metabolites database (4400 compounds) and HMDB Feces metabolites (v5. 6790 compounds). Features without MS2 or with a peak quality and QC peak quality of less than 5.5 were excluded from statistical analyses. An independent samples t-test was performed for statistical analysis, and feature significance was defined a *p*<0.05 and abs(log_2_(fc))>0.5.

## Data availability

Mouse metadata is available in supplementary file S1. Forced swim test video data are available at Zenodo (https://zenodo.org/) with DOI 10.5281/zenodo.21652181. The 16S rRNA gene amplicons supporting the findings of this study are deposited and available in NCBI Sequence Read Archive (https://www.ncbi.nlm.nih.gov/genbank/) under the BioProject ID PRJNA1504233 with reviewer link (https://dataview.ncbi.nlm.nih.gov/object/PRJNA1504233?reviewer=trpth4iiao3tb0ujmurucjomdi). Proteomic, metaproteomic, untargeted metabolomic, and targeted SCFA metabolomic raw data are available in the MassIVE Repository (https://massive.ucsd.edu/ProteoSAFe/static/massive.jsp) and accessible with accession number MSV000102649, password Wrightlabstress1, ftp://MSV000102649@massive-ftp.ucsd.edu. The authors confirm that all data supporting the findings of this study are contained and available within the main text or the supplementary information. Any additional information is available from the lead contact upon request.

## Code availability

All R codes used for statistical analysis and data visualizations, and the Compound Discoverer workflow file are available in the GitHub repository associated with this manuscript accessible at https://github.com/acwickham/Slusser_2026_Social-isolation.

## Acknowledgements

Funding for this research was provided through the James R. Schofield Biomedical Endowment to A.T.W. This study was supported by the use of Baylor University’s core facilities. The authors sincerely thank the staff of Baylor University’s vivarium, where animals were housed and maintained, for their guidance and assistance during animal care and experiments. Sequencing and immunological marker analyses were performed in Baylor University’s Molecular Biosciences Center with the assistance and expertise of Dr. Michelle Nemec whom we thank for her support. We gratefully acknowledge Baylor University’s Mass Spectrometry Center, maintained by Dr. Alejandro J. Ramirez and Dr. Chinthaka Seneviratne, for proteomic, metaproteomic, and metabolomic data collection and analysis. We acknowledge the computing resources provided by Baylor University’s Kodiak High-Performance Computing cluster, directed by Mike Hucheson. We acknowledge the use of BioRender in generating parts of certain figures.

## Author contributions

Conceptualization, A.C.S. & A.T.W.; investigation, A.C.S., T.R.R., L.A.H., M.M.H., D.T.L., & T.J.E.; software, T.R.R.; formal analysis, A.C.S., T.R.R; writing – original draft, A.C.S.; writing – review and editing, A.C.S., A.T.W., L.A.H., T.R.R., M.M.H., D.T.L., & T.J.E. All authors read and approved the final manuscript.

## Competing Interests

The authors have no competing interests to declare.

## Notes

https://massive.ucsd.edu/ProteoSAFe/static/massive.jsp

https://dataview.ncbi.nlm.nih.gov/object/PRJNA1504233?reviewer=trpth4iiao3tb0ujmurucjomdi

### Competing Interest Statement

The authors have declared no competing interest.

