## Supplemental Information for "Chronic stress remodels gut bacterial ecology and polysaccharide metabolism with neuroendocrine consequences despite unchanged behavior"

**Supplementary Figures**

**Figure S1.** Host responses to social isolation stress

**Figure S2.** Effects of host stress on large intestine taxonomy and luminal short chain fatty acid concentrations

**Figure S3.** Male metaproteomic analysis and significant stress response proteins

**Figure S4.** Female metaproteomic analysis and top genera based on significant protein proportion

**Figure S5.** Significantly altered carbohydrate-active enzymes in male and female metaproteomes

**Figure S6.** Circulating estrone-glucuronide levels, altered transcription factor activity and detected proteins in the estrogen signaling pathway

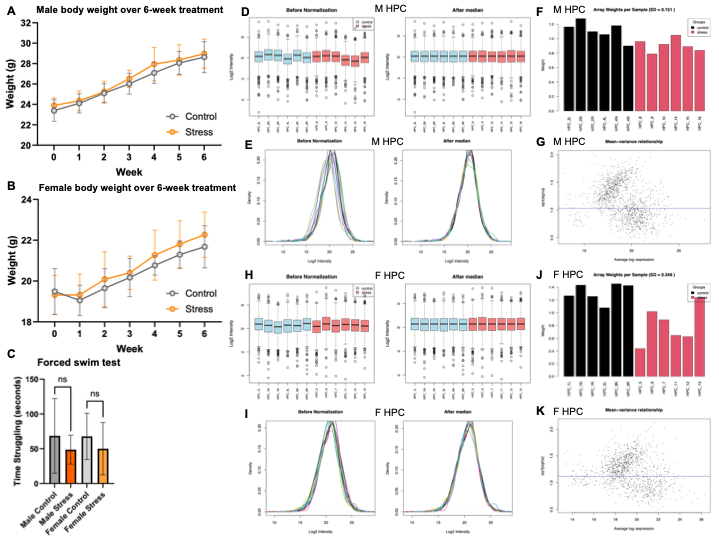

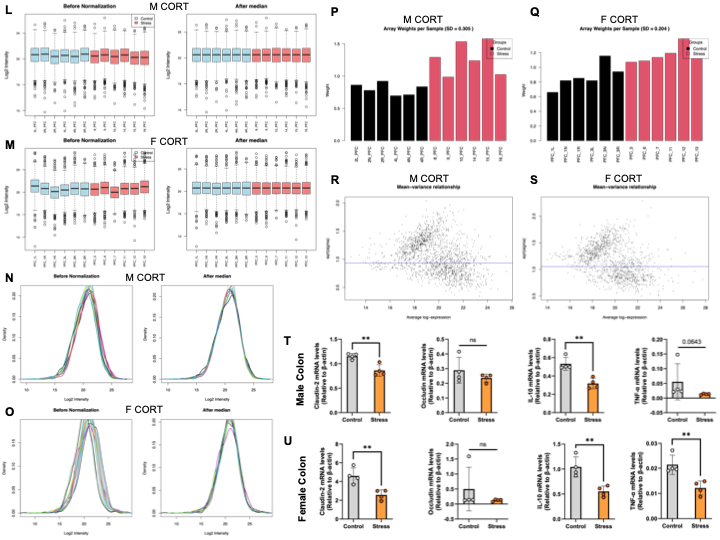

**Figure S1**. **Host responses to social isolation stress.** **A & B** Male and female weekly body weight measurement during the treatment period. Statistical significance was calculated using a two-sample t-test where significance as defined as *p*<0.05. Error bars represent standard deviation from the mean. **C** Forced swim test results. Statistical significance was calculated using a two-sample t-test where significance as defined as *p*<0.05. Error bars represent standard deviation from the mean. **D &** **E** Median normalization was performed across male hippocampus samples. Whiskers extend to the most extreme values within 1.5X the interquartile range. **F** Sample array weights before correction using the limma R package (version 3.66.0), male hippocampus. **G** Mean-variance relationship of the imputed data linearly modeled by limma, male hippocampus. **H & I** Median normalization was performed across female hippocampus samples. Whiskers extend to the most extreme values within 1.5X the interquartile range. **J** Sample array weights before correction using limma, female hippocampus. **K** Mean-variance relationship of the imputed data linearly modeled by limma, female hippocampus. **L-O** Median normalization was performed across all male and female cortex samples. **P & Q** Sample array weights before correction using limma, male and female cortex. **R & S** Mean-variance relationship of the imputed data linearly modeled by limma, male and female cortex. **T-U** Immune response and tight junction markers measured in the colon. Statistical significance was determined by independent samples t-test assuming equal variance, significance was defined as *p*<0.05: not significant (ns), **p*<0.05, ***p*<0.01. Error bars represent standard deviation from the mean.

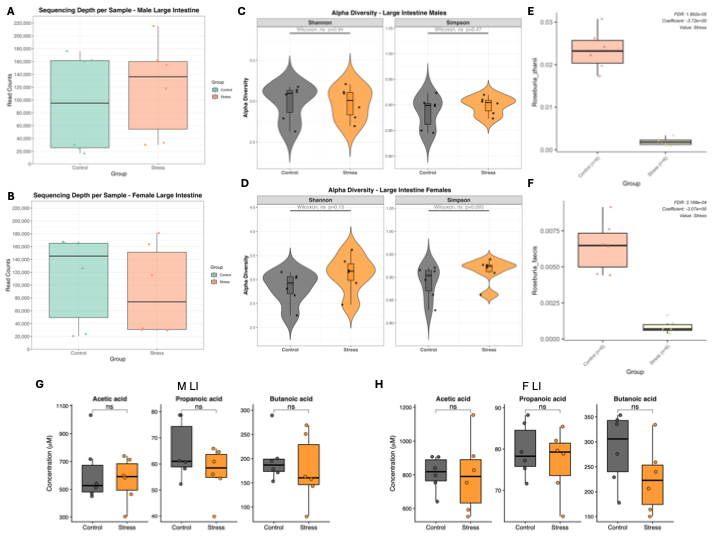

**Figure S2. Effects of host stress on large intestine taxonomy and luminal short chain fatty acid concentrations**. **A & B** Box plots showing 16S sequencing depth per sample of male and female large intestinal content. Whiskers extend to the most extreme values within 1.5X the interquartile range. **C & D** Alpha diversity (Shannon and Simpson diversity indices) of male and female large intestine. Wilcoxon rank-sum test was used to determine statistical significance, where *p*<0.05 was significantly different. Whiskers extend to the most extreme values within 1.5X the interquartile range. **E** Differential *Roseburia zhanii* species abundance in male LI. **F** Differential *Roseburia faecis* species abundance in male LI. Statistical significance was calculated using MaAsLin2 linear models. Raw p-values were adjusted using the Benjamini-Hochberg correction. Statistical significance was defined as *q*<0.05. Whiskers extend to the most extreme values within 1.5X the interquartile range. **G & H** Short chain fatty acid concentrations in the LI of male and female control and stress mice. An independent samples t-test assuming equal variance was used to determine statistical significance where *p*<0.05 was statistically different. Whiskers extend to the most extreme values within 1.5X the interquartile range.

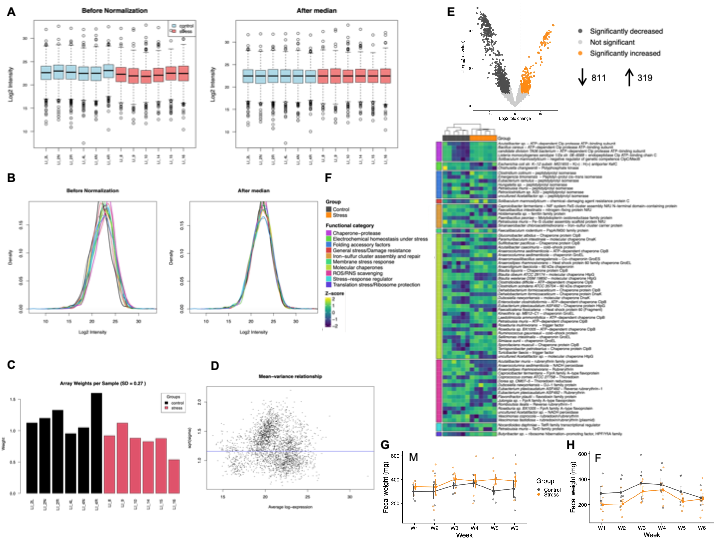

**Figure S3. Male metaproteomic analysis and significant stress response proteins. A & B** Median normalization was performed across all samples. **C** Sample array weights before correction using the limma R package (version 3.66.0). **D** Mean-variance relationship of the imputed data linearly modeled by limma. **E** Volcano plot showing differential expression of proteins identified in the male large intestine metaproteome. Statistical analysis was conducted using a weighted linear model and empirical-Bayes moderated t-test method through limma. Raw *p*-values were adjusted using Benjamini-Hochberg (BH) false discovery rate adjustment (*q*-value). Significance was defined as *q*<0.05 and an absolute log2 fold change ≥2. Statistical analysis revealed that 811 proteins were significantly down regulated while 319 proteins were significantly upregulated. **F** Heatmap of significantly altered bacterial stress response protein abundances. Color-coded functional categories are shown one the left. Z-score normalized intensities shown in purple (negative) to yellow (positive). **G & H** Fecal production analysis over study duration. Statistical significance was calculated using a two-sample t-test where significance as defined as *p*<0.05. Error bars represent standard error.

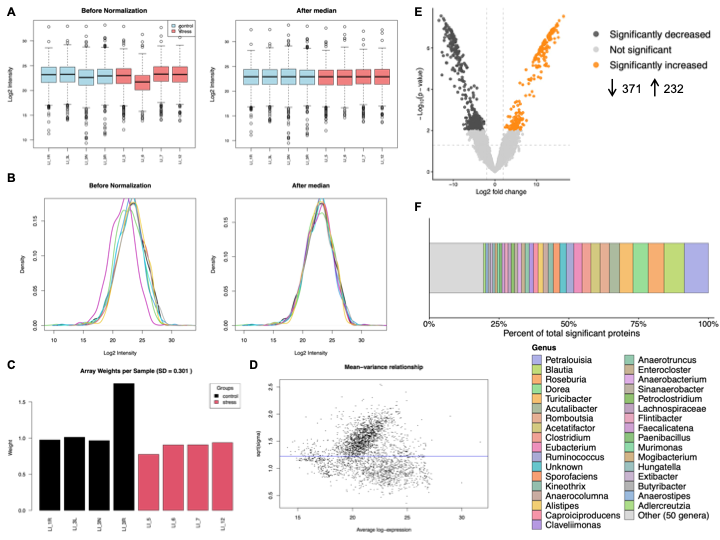

**Figure S4. Female metaproteomic analysis and top genera based on significant protein proportion. A & B** Median normalization was performed across all samples. **C** Sample array weights before correction using limma R package (version 3.66.0). **D** Mean-variance relationship of the imputed data linearly modeled by limma. **E** Volcano plot showing differential expression of proteins identified in the male large intestine metaproteome. Statistical analysis was conducted using a weighted linear model and empirical-Bayes moderated t-test method through limma. Raw *p*-values were adjusted using Benjamini-Hochberg (BH) false discovery rate adjustment (*q*-value). Significance was defined as *q*<0.05 and an absolute log2 fold change ≥2. Statistical analysis revealed that 371 proteins were significantly down regulated while 232 proteins were significantly upregulated. **F** Stacked bar chart showing the most abundant genera by percent of total significant proteins.

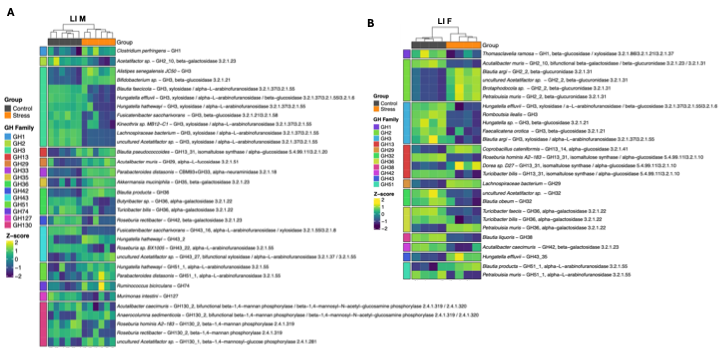

**Figure S5. Significantly altered carbohydrate-active enzymes in male and female metaproteomes. A & B** Significantly altered proteins in male mice microbiomes (n=6 per group) and female mice microbiomes (n=4 per group) with predicted glycoside hydrolase activity. Color-coded glycoside hydrolase family groupings shown on the left. Z-score normalized intensities shown in purple (negative) to yellow (positive).

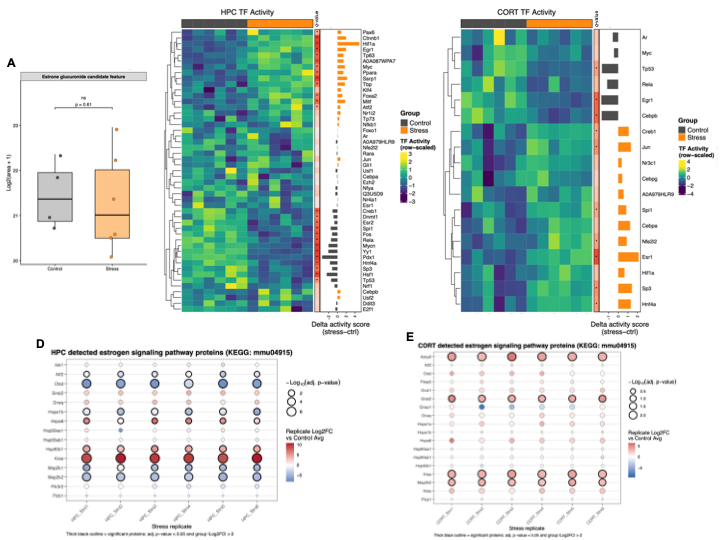

**Figure S6. Circulating estrone-glucuronide levels, altered transcription factor activity and detected proteins in the estrogen signaling pathway. A** Box plot showing circulating estrone glucuronide candidate levels in SI and GH females. Statistical significance determined by independent samples t-test assuming equal variance where a significant result is defined by *p*<0.05*.* Whiskers extend to the most extreme values within 1.5X the interquartile range. **B & C** Heatmaps of significantly altered transcription factor activity scores in female HPC and CORT regions, represented by z-scores. Statistical analyses were performed using a Welch’s t-test and *p-*value adjustment with Benjamini-Hochberg correction so that statistical significance was defined as *q*<0.05. The red-gradient bar on the right side of the heat map represents the intensity of the *q*-value, where darker shades of red are smaller, more significant values. Statistical significance is indicated by the asterisk. The horizontal bar chart on the far right from the heatmap represents the magnitude of the fold change in transcription factor activity scores between the groups. **D & E** Detected, annotated proteins belonging to KEGG pathway mmu04915 murine estrogen signaling in female HPC and CORT brain regions. Significant proteins are indicated by a black outline. Bubble fill color indicates protein expression fold change in SI compared to GH mice.
